# A human pluripotent stem cell-based somitogenesis model using microfluidics

**DOI:** 10.1101/2023.10.29.564399

**Authors:** Yue Liu, Yung Su Kim, Xufeng Xue, Norio Kobayashi, Shiyu Sun, Qiong Yang, Olivier Pourquié, Jianping Fu

## Abstract

Emerging human pluripotent stem cell (hPSC)-based embryo models are useful for studying human embryogenesis. Particularly, there are hPSC-based somitogenesis models using free-floating culture that recapitulate somite formation. Somitogenesis *in vivo* involves intricately orchestrated bio-chemical and -mechanical events. However, none of the current somitogenesis models controls biochemical gradients or biomechanical signals in the culture, limiting their applicability to untangle complex biochemical-biomechanical interactions that drive somitogenesis. Here we report a new human somitogenesis model by confining hPSC-derived presomitic mesoderm (PSM) tissues in microfabricated trenches. Exogenous microfluidic morphogen gradients imposed on PSM cause axial patterning and trigger spontaneous rostral-to-caudal somite formation. A mechanical theory is developed to explain the size dependency between somites and PSM. The microfluidic somitogenesis model is further exploited to reveal regulatory roles of cellular and tissue biomechanics in somite formation. This study presents a useful microengineered, hPSC-based model for understanding the bio-chemical and -mechanical events that guide somite formation.

## INTRODUCTION

The formation of morphological boundaries between developing tissues is an integral mechanism for generating body forms and functions. In particular, formation of somites dictates the body layout of a vertebrate embryo and ultimately the structure of its musculoskeletal system. During somitogenesis, the presomitic mesoderm (PSM), a bilateral strip of mesenchymal tissue flanking the forming neural tube, progressively segments into bilaterally symmetrical epithelial somites in a rostral (R)-to-caudal (C) direction (**Fig. 1a**). New somites are formed at the rostral end of the PSM, where cells undergo mesenchymal-epithelial transition (MET) and coalesce into a rosette-like structure that pulls apart from the PSM. Although theories have been proposed to correlate somitogenesis dynamics with biochemical signals such as antiparallel gradients of FGF / WNT / retinoic acid (RA) along the R-C axis in the PSM^1–4^, detailed mechanisms underlying somitogenesis remain elusive owing to the inaccessibility of *in vivo* models for modulating endogenous morphogen gradients, especially for mammalian species. Furthermore, despite that somites acquire their distinct morphology *via* intricately coordinated biomechanical events, the biomechanical driving force leading to somite formation and size regulation remains obscure.

**Figure 1.**
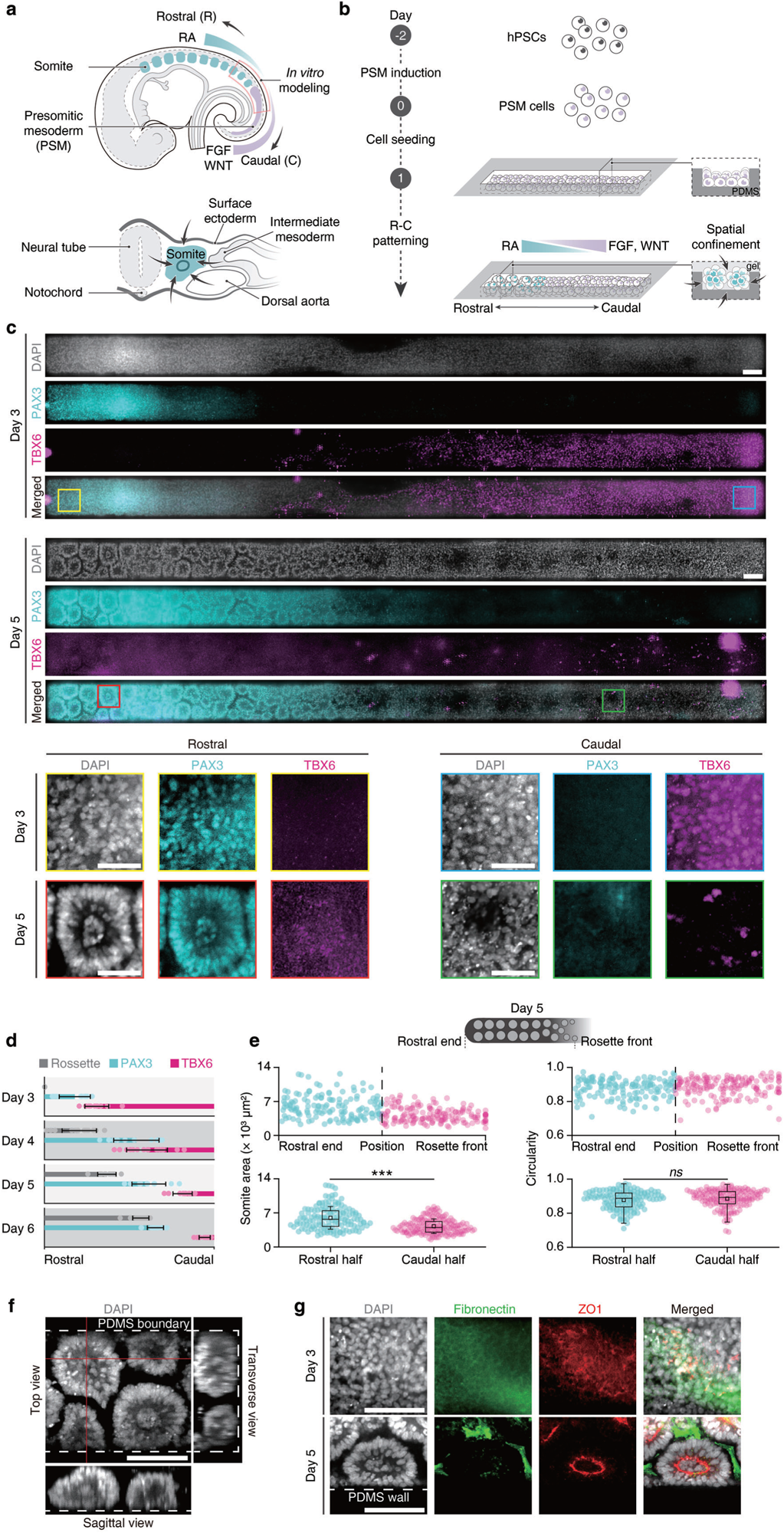
A hPSC-based, microfluidic somite development model (µSDM). **a**, (top) Sagittal view of a vertebrate embryo showing somite formation from the presomitic mesoderm (PSM), regulated by opposing morphogen gradients along the rostral (R)-caudal (C) axis. The µSDM is developed to model somite formation from the thoracic / lumbar PSM region, as marked in the dashed box. (Bottom) Transverse section of the trunk showing a somite (cyan) mechanically confined by adjacent tissues during its development. **b**, Schematics showing µSDM development protocol. See **Methods** for µSDM design considerations. hPSC-derived PSM tissues are confined in microfabricated open trenches, before 3D gel overlay and microfluidic morphogen gradients are imposed on PSM tissues to establish an R-C axis and drive spontaneous somite formation, beginning rostrally and extending caudally. **c**, Representative stitched confocal micrographs showing µSDM on Day 3 and Day 5 stained for PAX3 and TBX6. Cell nuclei were counterstained with DAPI. Zoom-in views of boxed regions are shown at the bottom. Scale bars, 100 µm for full-tissue images and 50 µm for zoom-in images. **d**, Bar plot showing spatial patterns of somite formation and PAX3 and TBX6 expression in µSDM as a function of culture day as indicated. *n*_µSDM_ ≥ 10 for each day, and data are plotted as the mean ± s.d. **e**, (top) Area and circularity of individual somites between the rostral end and rosette front in µSDM on Day 5. (bottom) Data from the rostral and caudal halves within this somite formation regime are grouped and analyzed respectively. *n*_µSDM_ = 12, and *n*_somite_ = 140 for rostral halves and *n*_somite_ = 163 for caudal halves. Boxes and bars indicate interquartile ranges and median values, respectively, and squares and error bars indicate the mean ± s.d. Two-sample *t*-tests for somite area comparison (*P* = 1.5×10^-13^) and for somite circularity comparison (*P* = 0.20). **f**, Representative confocal Z-stack images showing individual somites in µSDM on Day 5 stained with DAPI. Scale bar, 100 µm. **g**. Representative confocal micrographs showing rostral regions of µSDM on Day 3 and Day 5 stained for fibronectin and ZO1 as indicated. Scale bars, 100 µm.

The recent emergence of hPSC-based embryo models opens up exciting opportunities to promote fundamental understanding of human development^5–7^. Particularly, there are hPSC-based somitogenesis models recently developed based on three-dimensional, free-floating cultures that show somite formation^8–12^. However, none of the models controls biochemical gradients or biomechanical signals in the culture, limiting their applicability to untangle complex biochemical-biomechanical interactions that drive somitogenesis. In this work we report a new human somite formation model by mechanically confining hPSC-derived PSM tissues in microfabricated trenches to simulate an essential mechanical boundary condition for PSM development *in vivo*^13,14^. Controlled exogenous microfluidic morphogen gradients are then imposed on the PSM tissues to drive their R-C patterning and trigger spontaneous somite formation, beginning rostrally and extending caudally. Leveraging the compatibility of this human somite formation model with live imaging and biomechanical and molecular characterizations and perturbations, we further experimentally and theoretically explored the mechanical regulators that contribute to somite formation and size regulation.

## RESULTS

### A hPSC-based, microfluidic somite development model

To model the sculpting of somites, we developed a hPSC-based, microfluidic somite development model (µSDM). Specifically, µSDM develops in a polydimethylsiloxane (PDMS)-based microfluidic device containing three channels partitioned by circular support posts (see **Methods** for µSDM design considerations; **Supplementary Fig. 1a&b**). Rectangular micro-trench structures (width: 200 µm; length: 4 mm; depth: 200 µm) at the bottom surface of the central channel of the microfluidic device are used to position and contain hPSC-derived PSM tissues, to mimic their geometrical confinements and mechanical boundary condition *in vivo*^13,14^ (see **Methods**). With different signaling molecules supplemented to the two reservoirs of the central channel, gradients of developmental signals, such as RA, FGF and WNT, could be explicitly imposed on PSM cells seeded in micro-trenches through passive diffusion (**Supplementary Fig. 1a&c**). Indeed, passive diffusion assays using fluorescent dextran as a proxy in the central channel of the microfluidic device confirm the establishment of stable chemical gradients within about 36 h (**Supplementary Fig. 1c**). Specifically, to derive PSM cells, H9 human embryonic stem cells (hESCs) in tissue culture plates are treated with a basal medium (BM) supplemented with CHIR99021 (or CHIR, a WNT activator) and LDN 193189 (or LDN, a BMP inhibitor), referred herein to as CL medium, for two days (from Day -2 to Day 0; **Fig. 1b** and **Supplementary Fig. 1a&d**). On Day 0, hESCs differentiate into PSM cells expressing TBX6 (PSM-specific marker) but not SOX2 (pluripotency maker) or PAX3 (somite marker), with TBX6^+^ PSM cell proportion around 94.4% (**Supplementary Fig. 1d**). TBX6^+^ PSM cells co-express HOXC9 but not HOXC10, suggesting their thoracic or rostral lumbar axial identity (**Supplementary Fig. 1d**). PSM cells are collected from tissue culture plates before re-seeding at a density of 15 × 10^6^ cells mL^-1^ into the central channel of the microfluidic device on Day 0 in CL medium. Since only interior walls and bottom surfaces of micro-trenches are coated with Geltrex and thus adhesive to cells, whereas the other areas in the central channel are coated with bovine serum albumin (see **Methods**; **Supplementary Fig. 1a**), PSM cells loaded into the central channel are deposited only into micro-trenches and form initial PSM tissues (**Supplementary Fig. 1e**). On Day 1, Geltrex is loaded into the central channel to establish a Geltrex overlay for PSM tissues. Simultaneously, PD 173074 (or PD, a FGFR inhibitor) and RA are supplemented into the left reservoir of the central channel, hereafter designated as the rostral (R) end, while CHIR, LDN and FGF8 are added to the right reservoir, designated as the caudal (C) end (**Supplementary Fig. 1a**). This way, R-to-C gradient of RA and C-to-R gradients of FGF and WNT signals are established over the length of PSM tissues, mimicking morphogen environments experienced by PSM tissues *in vivo*.

Based on brightfield imaging of µSDM development, boundaries splitting PSM tissues into small compartments become visible in rostral µSDM regions from Day 4 onwards and gradually propagate caudally over time (**Supplementary Fig. 1e**). On Day 6, well-defined tissue boundaries separating individual rosette-like structures, indicating successful somite formation, are clearly notable in the rostral half of µSDM (**Supplementary Fig. 1e**). Immunostaining was conducted for TBX6 and PAX3 on µSDM (**Fig. 1c** and **Supplementary Fig. 1g**). On Day 3, R-C patterned expression of TBX6 and PAX3 in µSDM emerges, with cells in the most rostral region expressing PAX3 but not TBX6 (**Fig. 1c**). PAX3^+^ somitic domain in µSDM expands caudally between Day 3 and Day 6, with concurrent caudal regression of TBX6*^+^* PSM region with a comparable speed (**Fig. 1c&d**). Development of PAX3^+^ somitic domain precedes somite formation in µSDM (**Fig. 1c&d**). Nonetheless, on Day 3 local cellular compaction and re-organization become evident in rostral µSDM regions (**Fig. 1c**), indicating initiation of MET and somite formation. On Day 6, well separated, mature PAX3^+^ somites with an epithelial appearance and a closely packed circumferential ring of columnar-shaped cells surrounding small clumps of somitocoel cells are detectable across the entire rostral half of µSDM (**Fig. 1c&d** and **Supplementary Fig. 1g**). Immunostaining of µSDM on Day 5 for HOXC9 and HOXC10 shows uniform HOXC9 expression yet with a few HOXC10^+^ cells, confirming the thoracic or rostral lumbar axial identity of µSDM (**Supplementary Fig. 1h**).

We next examined gene expression pattern in µSDM by dissecting µSDM into three equal-length segments and collecting them for RT-qPCR (**Supplementary Fig. 1i&j**). Consistent with immunostaining, rostral µSDM segments show greater expression of somite-related markers *PAX3*, *FOXC2*, *MEOX1* and *TCF15*, whereas middle and caudal µSDM segments exhibit higher expression of PSM-related genes *HES7* and *TBX6* (**Supplementary Fig. 1i,j**).

To quantify spatiotemporal dynamics of somite formation, all somites in µSDM on Day 5 were split into two groups based on their relative distances to the caudal rosette formation front (**Fig. 1e**). Somites in the rostral half of the rosette-forming domain have an average area of 5,973 µm^2^ (effective diameter of 87.2 µm; **Fig. 1e**). Somites in the caudal half of the domain, which include relatively newly formed somites, show a smaller average area of 4,257 µm^2^ (effective diameter of 73.6 µm; **Fig. 1e**). Somite sizes in µSDM are thus comparable with those in CS9-10 human embryos^8,9,11^. Circularities of somites in µSDM on Day 5 are comparable between the rostral and caudal halves of the rosette-forming domain (**Fig. 1e**). Thus, our data suggest that nascent somites in µSDM experiences a growth process that increases their sizes but maintains their shape.

Confocal imaging of Day 5 µSDM confirms distant separations of neighboring somites and radial orientations of epithelialized somite boundary cells (**Fig. 1f**). *In vivo*, nascent boundaries between forming somites and the PSM are stabilized by epithelialization of somite boundary cells and assembly of extracellular matrix proteins in between^15^. Thus, we conducted immunostaining of µSDM for fibronectin and ZO1 on Day 3 and Day 5. There is no clear spatial pattern of fibronectin or ZO1 in µSDM on Day 3 (**Fig. 1g**). On day 5, however, assembly of fibronectin is notable between adjacent somites, and ZO1 is evident demarcating the inner surface of epithelialized somite boundary cells, supporting the establishment of apical-basal polarity in individual somites (**Fig. 1g**).

Somite formation in µSDM is highly efficient and reproducible (**Supplementary Fig. 1f**), with about 93.8% samples successfully showing epithelialized rosette structures with well-separated tissue boundaries at the rostral ends.

To examine the robustness of µSDM, µSDM protocols were repeated using H1 hESCs and a hiPSC line (see **Methods**). Both H1 hESCs and the hiPSC line generate µSDM with R-C patterned TBX6 and PAX3 expression and well separated, PAX3^+^ somites at the rostral ends (**Supplementary Fig. 2a**). Progression of somite formation front is comparable between H9 and H1 hESCs and the hiPSCs (**Supplementary Fig. 2b**). Areas of somites in µSDM generated from H9 and H1 hESCs appear to be greater than those from the hiPSC line (**Supplementary Fig. 2c**).

We also tested different cell seeding densities ranging from 8 × 10^6^ cells mL^-1^ to 25×10^6^ cells mL^-1^ for H9 hESCs (**Supplementary Fig. 2d**), with data showing little contrast in somite formation propagation speed and somite area in µSDM (**Supplementary Fig. 2e&f**), indicating saturation of cell seeding density in µSDM protocols. Consistently, the thickness of PSM cell layers remaining in micro-trenches on Day 1 appears similar across the three tested cell seeding conditions (**Supplementary Fig. 2d**). Another mechanical factor, micro-trench width, was also explored (**Supplementary Fig. 2g**). Similar somite area and propagation speed are observed in µSDM generated from 200 µm-, 300 µm- and 400 µm-wide trenches (**Supplementary Fig. 2h&i**). The 200 µm-wide trenches, which were used in the rest of this study, give rises to approximately 2 rows of somites, whereas wider trenches produce more rows (**Supplementary Fig. 2j**). If somite dimension along the R-C axis of the PSM is dictated by extrinsic factors, such as morphogen gradients and forming somite-PSM interactions (more discussions below), we conjecture that the lateral dimension of somites is regulated by intrinsic factors such as tissue surface tension, which prefers a dimensional aspect ratio of unity and leads to similar somite areas across different micro-trench widths. External FGF8 gradients was also modulated for µSDM development (**Supplementary Fig. 2k**). Shallower FGF8 gradients result in expedited somite formation propagation (**Supplementary Fig. 2l**), even though somite areas appear insensitive to FGF8 gradient magnitudes tested (**Supplementary Fig. 2m**).

As shown in **Supplementary Fig. 1e**, µSDM is compatible with live imaging. To exploit this advantage, we further conducted brightfield live imaging of developing µSDM (**Supplementary Fig. 5a** and **Supplementary Video 1**). We identified individual somites at the caudal µSDM region and tracked their development from Day 4 to Day 6 (**Supplementary Fig. 2n-p**). Time-lapse analysis reveals that nascent somites in caudal µSDM regions adopt a small area before growing between Day 4 to Day 6, while circularity of the somites maintains a relatively constant value throughout this time (**Supplementary Fig. 2n-p**), consistent with data in **Fig. 1e**. We further employed a H9-H2B reporter line and tracked cellular dynamics in a forming somite at the rosette formation front *via* confocal imaging. MET-induced cellular compaction and epithelization are clearly evident in the forming somite, leading to a reduced in-plane tissue width and radially reoriented somitic cells setting up its boundary from adjacent somites (**Supplementary Fig. 2q** and **Supplementary Video 2**). Growth of nascent somites following their epithelialization was also observed, with dimensions of both somite and somitocoel increasing over time, likely involving active cell division within the epithelial ring and its lumen expansion (**Supplementary Fig. 2r** and **Supplementary Video 3**). Interestingly, somitic cells in the epithelial ring constantly delaminate and move towards somitocoel cells as either single cells or cell clusters (**Supplementary Fig. 2s** and **Supplementary Video 4**), supporting continuous remodeling and dynamic cell movements within somites.

### Single-cell RNA-sequencing analysis

To investigate dynamics of µSDM development at the transcriptome level, single-cell RNA-sequencing (scRNA-seq) was conducted for Day 2, Day 3, Day 4 and Day 6 µSDM, respectively (**Fig. 2a**). Uniform manifold approximation and projection (UMAP) dimension reduction was conducted for integrated scRNA-seq dataset combining data from all four time points, revealing five distinct mesodermal cell clusters annotated as ‘caudal PSM / cPSM’, ‘rostral PSM / rPSM’, ‘nascent somite / N-SM’, ‘early somite / E-SM’ and ‘somite / SM’ based on lineage marker expression patterns (**Fig. 2b-d** and **Supplementary Fig. 3a&b**). A small neuronal cell lineage cluster was also identified (not shown); it was excluded from further analysis for clarity. We speculate that PSM cells seeded on Day 0 might contain a small number of neuromesodermal progenitors which can give rise to neuronal lineages. Most cells in Day 2 µSDM are annotated as cPSM cells (**Fig. 2e**). On Day 3, majority of cells in µSDM are identified as either rPSM or N-SM cells (**Fig. 2e**). Most cells in Day 4 µSDM switch to E-SM identity before progressing to SM fate on Day 5 (**Fig. 2e**). Both cPSM and rPSM clusters show upregulated expression of PSM-related genes such as *TBX6*, *MSGN1*, *RSPO3*, *DLL1* and *HES7* (**Fig. 2f** and **Supplementary Fig. 3a&b**). The rPSM cluster also shows upregulated expression of rostral PSM markers *MESP2* and *RIPPLY2* (**Fig. 2f** and **Supplementary Fig. 3a&b**). The somitic clusters N-SM, E-SM and SM all exhibit elevated expression of somitic markers *TCF15* and *PAX3* (**Fig. 2f** and **Supplementary Fig. 3a&b**). N-SM cluster also shows pronounced expression of *RIPPLY1*, while E-SM cluster is marked by greater expression of caudal somite/rostral PSM markers such as *FOXC2* compared to that of the SM cluster (**Fig. 2f** and **Supplementary Fig. 3a&b**). Expression of *HOX*5-9 genes, but not more caudal *HOX* genes, is evident in µSDM between Day 2 and Day 6 (**Supplementary Fig. 3c**), consistent with immunostaining data in **Supplementary Fig. 1h**. Consistent with other *in vitro* human somitogenesis models^11^, RA signaling-attenuating gene *DHRS3* is upregulated in cPSM, whereas RA synthesis-associated genes *ALDH1A2* and *RDH10* show elevated expression in somitic clusters N-SM, E-SM and SM (**Supplementary Fig. 3d**). *FGF3*/*8*/*17*/*18* and *DUSP6*, an FGF signaling target gene, mainly express in cPSM and rPSM (**Supplementary Fig. 3d**).

**Figure 2.**
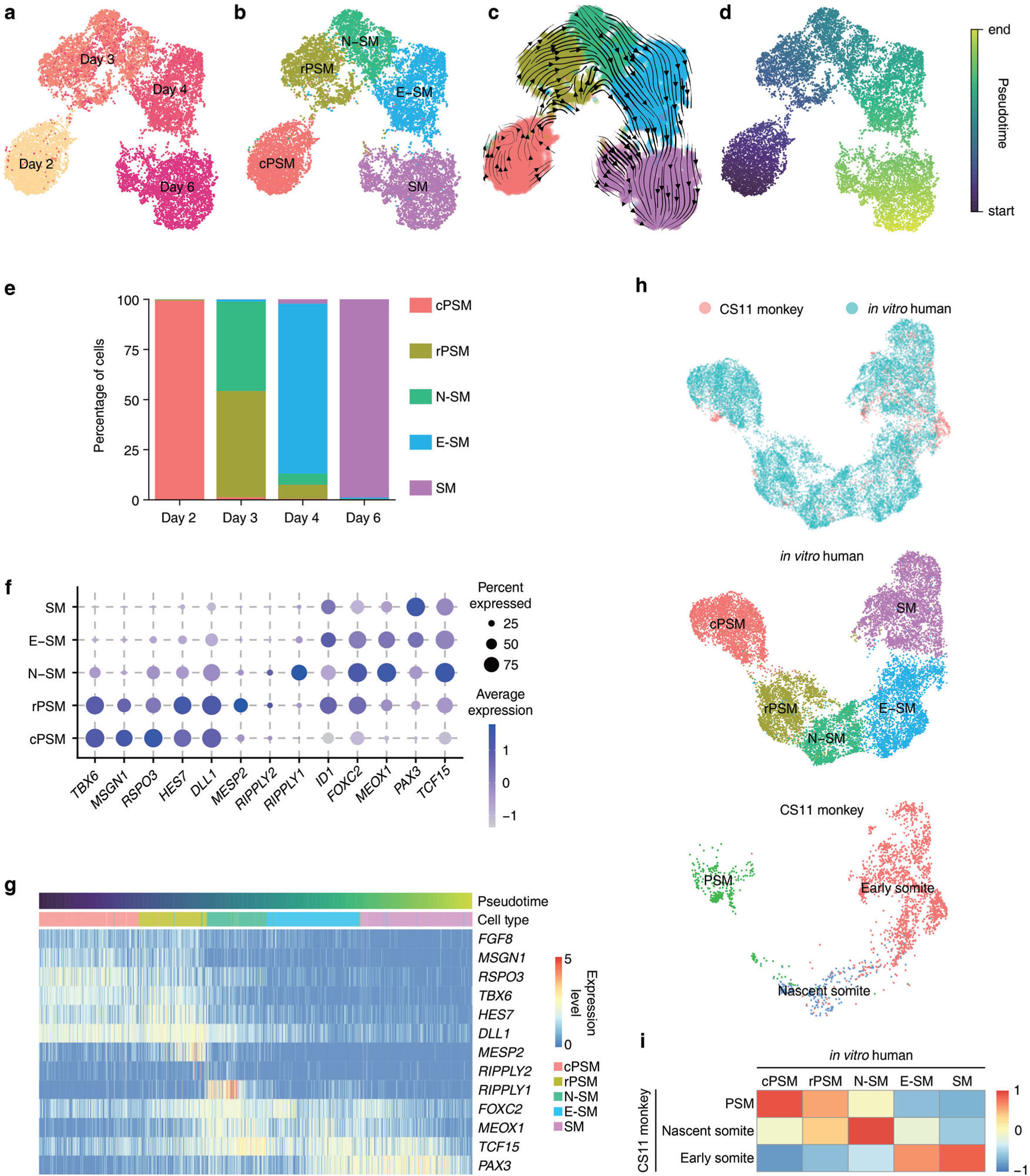
Single-cell transcriptomic analysis of µSDM. **a-d**, UMAP embedding of integrated single-cell transcriptome dataset of µSDM on Day 2 (*n*_cell_ = 3,259), Day 3 (*n*_cell_ = 3,808), Day 4 (*n*_cell_ = 3,620) and Day 6 (*n*_cell_ = 3,634), color-coded by µSDM culture time (**a**), cell identity annotation (**b&c**) and pseudotime (**d**). RNA velocity vectors projected onto UMAP embeddings in **c** show major cell progression directions in transcriptional space. Start and endpoints of arrows indicate observed-current and predicted-future cell states, respectively. **e**, Proportions of different cell types in µSDM over time. **f**, Dot plot showing expression of key marker genes across different cell clusters in µSDM. Dot sizes and colors indicate proportions of cells expressing corresponding genes and their averaged scaled values of log-transformed expression, respectively. **g,** Expression dynamics of key marker genes in µSDM along the pseudotime trajectory corresponding to **d**. Color bars above the heat map indicate pseudotime and cell identity as indicated. **h,** (top) UMAP projection of integrated scRNA-seq dataset from µSDM and somitogenesis-related cells from a CS11 monkey embryo^16^. (middle/bottom) UMAP projections of datasets from µSDM and the CS11 monkey embryo, separated from the integrated UMAP plot on the top, with cell identity annotations indicated. **i,** Pearson’s correlation analysis of µSDM cell clusters with somitogenesis-related cell clusters in the CS11 monkey embryo. Correlation coefficients between indicated µSDM and monkey cell clusters are calculated based on variable genes identified from the µSDM/monkey somitogenesis-related cell clusters (**Supplementary Table 2**). cPSM, caudal PSM; rPSM, rostral PSM; N-SM, nascent somite; E-SM, early somite; SM, somite.

Cell clustering analysis using UMAP in **Fig. 2a&b** shows cell fate transitions in µSDM from cPSM to rPSM and then somitic fates. Consistently, RNA velocity and pseudo-time analyses confirm a PSM-to-somite cell fate transition developmental trajectory (**Fig. 2c&d**). Along the pseudo-time trajectory, PSM markers *TBX6*, *MSGN1* and *RSPO3* are gradually down-regulated, whereas somitic markers *MEOX1*, *TCF15* and *PAX3* are up-regulated (**Supplementary Fig. 3e**). Some rostral PSM and early somite markers, such as *MESP2* and *RIPPLY1*/*2*, are transiently up-regulated in rPSM and N-SM cells before decreasing rapidly in E-SM and SM cells (**Fig. 2g** and **Supplementary Fig. 3e**).

We also conducted comparative transcriptome analysis, using scRNA-seq data from CS11 *cynomolgus* (*cy*) monkey embryo as a reference^16^. This comparative analysis reveals a reasonable overlap in UMAP projection between µSDM cells and cells from *cy* monkey PSM, nascent somite, and early somite clusters (**Fig. 2h**). On the basis of most variable genes identified from the integrated scRNA-seq dataset combining µSDM and *cy* monkey cells, cPSM/rPSM, N-SM and E-SM/SM clusters in µSDM show the closest correlations with *cy* monkey PSM, nascent somite and early somite cells, respectively (**Fig. 2i**). In addition, expression profiles of essential PSM and somitic markers in UMAP analyses show close similarities between µSDM cells and corresponding *cy* monkey somitogenesis-related cells (**Supplementary Fig. 3f**).

### HES7 dynamics along the R-C axis

Somitogenesis *in vivo* is a rhythmic process that correlates temporally with a molecular oscillator, or the segmentation clock, acting in PSM cells^17^. The segmentation clock drives dynamic and periodic expression of a number of so-called ‘clock’ genes, including *HES7*, across the PSM in a C-to-R fashion (**Fig. 3a**). To examine dynamic activities of clock genes in µSDM, a HES7 reporter hESC line^18^ was used for µSDM development and imaged with confocal microscopy continuously between Day 2 and Day 4 (**Fig. 3b&c** and **Supplementary Video 5**). Initially, HES7 traveling waves occur at both the rostral and caudal ends of µSDM, moving towards tissue center (**Fig. 3b&c**). This observation is comparable to edge-to-center traveling waves noted in other somite organoids without R-C axial patterning^10^. After initial few oscillations, rostral-to-center HES7 traveling waves dim down by Day 3 (**Fig. 3b&c**). The C-to-R HES7 traveling waves, however, persist, consistent with the establishment of an R-C axis in µSDM by Day 3 (**Fig. 3b&c**). To corroborate this finding, we analyzed mean HES7 intensities in rostral, middle and caudal regions of µSDM (**Fig. 3d**). HES7 oscillation in rostral µSDM regions flattens around Day 3, whereas HES7 intensities in middle and caudal µSDM regions continue undulating (**Fig. 3d**). To quantitatively analyze C-to-R HES7 traveling waves, we extracted their time intervals and traveling speeds (**Fig. 3b,e,f**). HES7 oscillation periods at the caudal µSDM end start from about 5 h, comparable with the period of human somite formation^18–20^, and increases to about 6.8 h on the fifth observed traveling wave (**Fig. 3e**). In contrast, HES7 oscillations at the caudal one-third position are notably slower, with its period starting from around 6.7 h and increasing up to 7 - 8 h (**Fig. 3e**). The difference of oscillation clock periods between the two µSDM locations is consistent with decreasing HES7 traveling wave speeds over time (**Fig. 3f**). The increasing oscillation clock periods at caudal µSDM locations are consistent with arrested segmentation clock when PSM cells in caudal µSDM regions transitioning to a somitic fate. The relatively shorter oscillation clock period at the µSDM caudal end is consistent with caudal-most cells in µSDM comparatively closer to the PSM identity. The spatial trend of increasing oscillation clock period in µSDM along its C-to-R axis is also consistent with observations in animal^21^ and *in vitro* models^12^.

**Figure 3.**
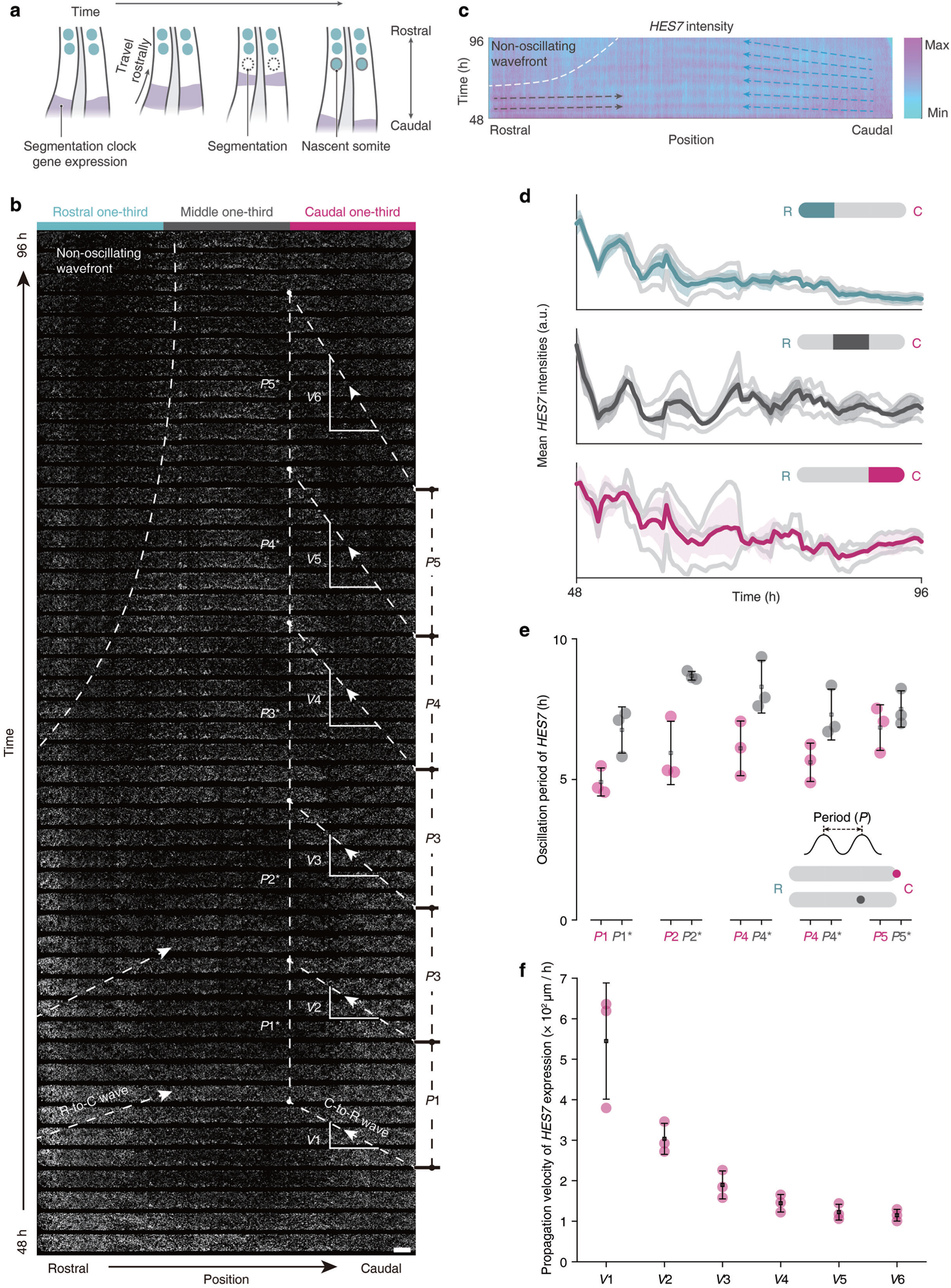
*HES7* dynamics during µSDM development. **a**, Schematics of oscillation and propagation of *HES7* expression in the PSM during somitogenesis *in vivo*. **b**, Time-lapse images showing development of µSDM using a *HES7* reporter line between *t* = 48 - 96 h. A dashed curve marks a non-oscillating region at the rostral end of µSDM. Traveling waves of *HES7* expression are marked by dashed lines with arrows. Oscillation periods at the caudal end (*P*1 - *P*5) and the caudal one-third point (*P*1* - *P*5*), and the wave speed between the two points (*V*1-*V*6) are marked and defined, respectively. Scale bar, 200 µm. **c**, Heatmap showing *HES7* intensity along µSDM length over time. *HES7* intensity is averaged across the micro-trench width. **d**, Mean *HES7* expression within the rostral, middle, and caudal one-third of µSDM as a function of time. Light grey lines indicate *HES7* expression dynamics of individual µSDM while colored lines represent averages. *n*_µSDM_ = 3, and shaded areas indicate s.d. **e**, Oscillation period of *HES7* at the caudal end (*P*1 - *P*5) and the caudal one-third point (*P*1* - *P*5*). Data are plotted as the mean ± s.d., with *n*_µSDM_ = 3. **f**, Propagation velocity of *HES7* expression waves traveling from the caudal end to the caudal one-third point. Data are plotted as the mean ± s.d., with *n*_µSDM_ = 3.

### Biomechanics regulates somite formation

We next utilized µSDM to explore the mechanical regulators that contribute to somite formation and size regulation. To this end, we developed a theoretical model based on the most essential mechanical factors involved in somite formation. At the rostral end of the PSM where cells transition to somitic fates, the cells undergo MET and coalesce into a rosette-like structure that pulls apart from the PSM^22–26^. Somite-forming cell clusters at the rostral end of the PSM spontaneously become more compact, inducing a contractile eigen-strain (an inelastic deformation) on the PSM and producing strain energy. When the strain energy exceeds surface energy required for the formation of a new somite-PSM boundary, the forming somite will delaminate from the rostral PSM and become a nascent somite (**Fig. 4a**). As the length of PSM *in vivo* needs to be compatible with and thus constrained by adjacent tissues in the trunk, it is reasonable to assume that the total length of PSM is fixed during the formation of a new somite, which therefore provides a boundary condition for PSM deformation. Through an energetic analysis (see **Methods**), a scaling law that connects the dimension of nascent somite (*d*) with the length of PSM (*L*) is acquired as *d* / *λ* = (*L* / *λ*)^1/2^, in which *λ* is a fitting parameter defined as *λ* = 4*γ* / *Eε*^*2^, with *γ* being tissue surface energy density, *E* PSM tissue stiffness, and *ε*\* eigen-strain. Despite its simplistic nature, this scaling law agrees reasonably well with *in vivo* data from mouse^27,28^, chick^28,29^, and zebrafish^28,30^ embryos (**Fig. 4b**). Furthermore, the scaling law also fits reasonably well with data generated from µSDM, as well as data from human embryo^31^ (**Fig. 4b**).

**Figure 4.**
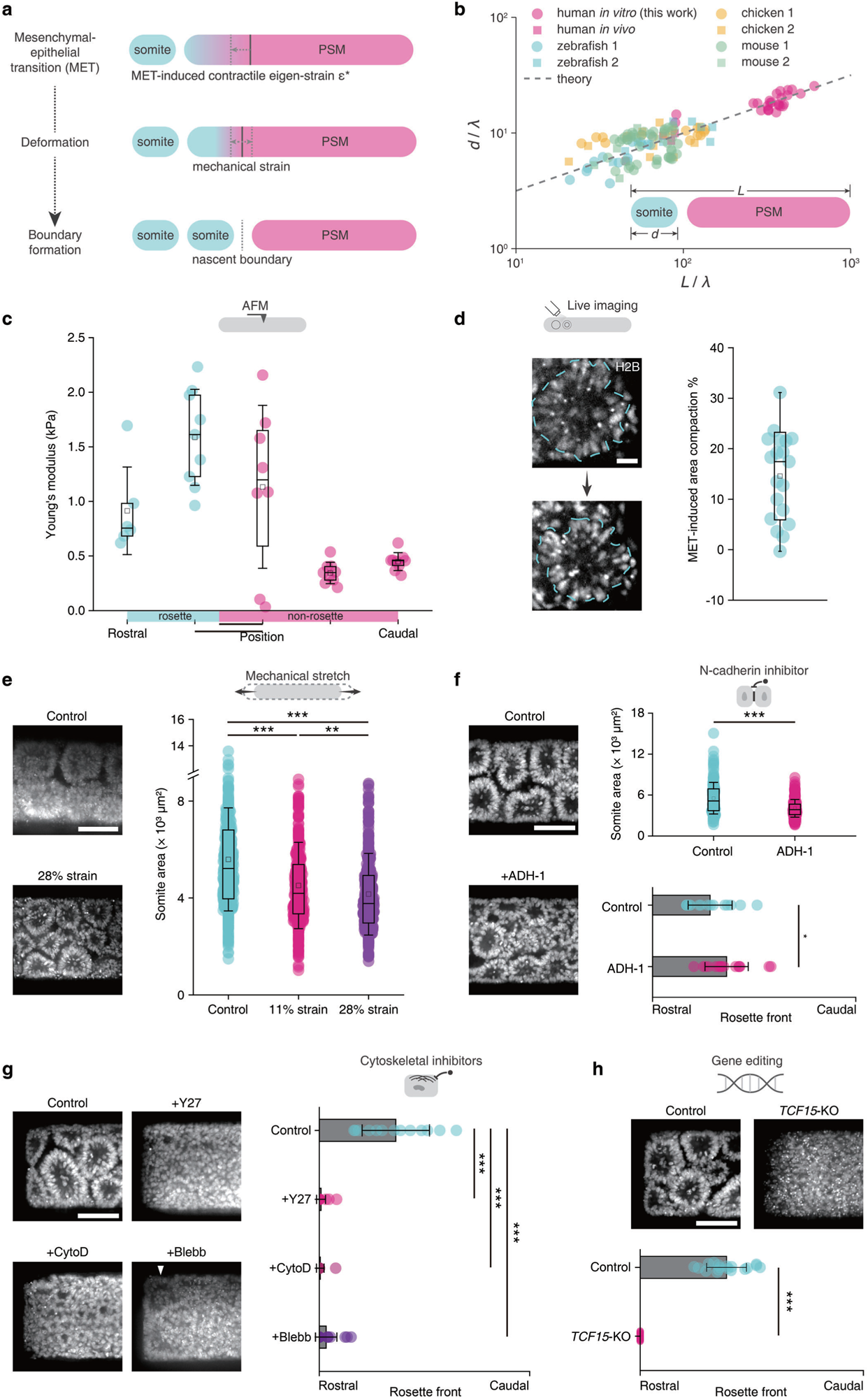
Mechanical modeling, characterization and perturbation of µSDM development. **a**, Schematics of MET-driven tissue deformation and nascent tissue boundary formation at the rostral PSM, leading to a forming somite delaminating from the PSM. **b**, Scaling law between length scales of a newly formed somite and the PSM derived from mechanical modeling, fitted with *in vivo* data from the zebrafish mouse^27,28^, chick^28,29^, zebrafish^28,30^ and human embryos,^31^ and µSDM data. Data from µSDM were obtained from *n*_µSDM_ = 26 on Day 4, 5 and 6. **c**, *Young*’s modulus measured by atomic force microscopy (AFM) for µSDM regions with and without rosette formation as indicated on Day 5. *n*_µSDM_ = 2. **d**, (left) Live imaging of µSDM development from a H2B reporter line showing compaction of a single forming somite between Day 4 and 5. Scale bar, 20 µm. (right) Quantitative data showing projected area compaction of single forming somites in µSDM. *n*_µSDM_ = 5. **e**, (left) Representative confocal micrographs showing µSDM on Day 6 stained with DAPI under control and 28% strain conditions. (right) Quantitative data showing projected area of individual somites in µSDM as a function of mechanical strains. Two-sample *t*-tests for control *vs.* 11% strain (*P* = 8.8×10^-14^), control *vs.* 28% strain (*P* = 1.3×10^-25^), and 11% strain *vs.* 28% strain (*P* = 6.4×10^-3^). In control group, *n*_µSDM_ = 36 and *n*_somite_= 507; in 11% strain group, *n*_µSDM_ = 18 and *n*_somite_ = 326; in 28% strain group, *n*_µSDM_ = 18 and *n*_somite_ = 376. **f**, Impact of ADH-1 treatment on µSDM development. (left) Representative confocal micrographs showing µSDM on Day 5 stained with DAPI under control and ADH-1 treatment conditions. (right) Quantitative data showing projected somite area and rosette formation front in µSDM on Day 5 under control and ADH-1 treatment conditions as indicated. For somite projected area data, *n*_µSDM_ = 14 and *n*_somite_ = 171 for control group and *n*_µSDM_ = 13 and *n*_somite_ = 207 for ADH-1 treatment group. Two-sample *t*-test for the two groups (*P* = 3.8×10^-14^). For rosette front data, *n*_µSDM_ = 15 for control group and *n*_µSDM_ = 16 for ADH-1 treatment group. Two-sample *t*-test for the two groups (*P* = 0.029). Somite area data in **e** and **f** were measured from full tissues. **g**, Impact of Y-27632 (Y27), cytochalasin D (CytoD) and blebbistatin (Blebb) treatment on µSDM development. (left) Representative confocal micrographs showing rostral regions of µSDM on Day 5 stained with DAPI under different conditions as indicated. While arrowhead marks occasional rosette-like cell organization under Blebb treatment. (right) Quantitative data showing rosette formation front in µSDM on Day 5 under different conditions as indicated. *n*_µSDM_ = 13 (control), *n*_µSDM_ = 20 (Y27), *n*_µSDM_ = 18 (CytoD), and *n*_µSDM_ = 18 (Blebb). Two-sample *t*-tests: control *vs.* Y27, *P* = 4.2×10^-11^; control *vs.* CytoD, *P* = 2.1×10^-10^; control *vs.* Blebb, *P* = 4.0×10^-9^. **h**, Effect of *TCF15*-KO on µSDM development. (top) Representative confocal micrographs showing µSDM on Day 5 stained with DAPI under different conditions as indicated. (bottom) Quantitative data showing rosette formation front in µSDM on Day 5 under different conditions as indicated. *n*_µSDM_ = 18 for both control and *TCF15*-KO conditions. Two-sample *t*-test between control vs. *TCF15*-KO conditions (*P* < 0.001). In **c**, **d** and all somite area quantification plots in **e**-**h**, boxes and bars indicate interquartile ranges and median values, respectively, and squares and error bars indicate the mean ± s.d. In rosette front quantification plots, bars and error bars indicate the mean ± s.d. Scale bars in **e**-**h**, 100 µm.

To examine fidelity of the scaling law, biomechanical characterizations were conducted on µSDM. PSM tissue stiffness *E* in µSDM was determined using atomic force microscopy (**Fig. 4c**). Eigen-strain *ε*\* was obtained by live imaging of the H2B reporter to record compaction of forming somites (**Fig. 4d**). Together with the *λ* used for best fitting of the scaling law with available µSDM somite size and PSM tissue length data, the surface energy density of PSM tissue was determined as *γ* = 4.4 pN µm^-1^, which is of a comparable order of magnitude with that measured for zebrafish mesoderm^32^ γ = 20.5 pN µm^-1^.

We next conducted biomechanical, biochemical and genetic perturbation assays to examine how somite formation is regulated mechanically. A cell-stretching device was employed to apply periodic tensile straining with a period of 4 h on µSDM between Day 5 and Day 6 (see **Methods**; **Supplementary Fig. 4a-d**). Somite area in µSDM reduces to 80.7% of control values at a peak strain of 11% and further down to 74.3% under a peak strain of 28% (**Fig. 4e**). When ADH-1, an N-cadherin inhibitor, was supplemented into µSDM culture, which should reduce intercellular surface energy and thereby facilitate somite boundary formation, somite area in µSDM decreases by 26.9% compared to untreated controls, together with a longer rosette-forming rostral region in µSDM (**Fig. 4f**). We next applied Y-27632 (ROCK inhibitor), cytochalasin D (actin polymerization inhibitor) and blebbistatin (myosin inhibitor) to inhibit intracellular cytoskeleton contraction machineries. Somite formation in µSDM is largely abolished under these drug inhibition conditions (**Fig. 4g**). Finally, we applied CRISPR-Cas9 gene editing tools to knockout (KO) *TCF15*, a gene involved in regulation of somitogenetic MET^33,34^, in H9 hESCs (**Supplementary Fig. 4e**). In *TCF15*-KO mouse embryo, epithelialization of PSM tissue and thus somite boundary formation are disrupted, leading to musculoskeletal patterning defects^33,34^. Consistent with *in vivo* murine phenotype, *TCF15*-KO completely inhibits epithelialization and somite boundary formation in µSDM (**Fig. 4h**).

## DISCUSSION

Despite its importance in defining the segmented body plan in vertebrate species, it remains challenging to study somitogenesis. Stem cell-based somitogenesis models are promising for advancing fundamental understanding of somitogenesis. However, existing hPSC-derived somitogenesis models lack extrinsic controls of bio-chemical or -mechanical cues, two essential mediators of somite formation, and as such they remain suboptimal for disentangling biochemical-biomechanical interactions that drive the sculpting of somites. The µSDM utilizes microfluidic morphogen gradients and microfabricated cell culture surfaces to effectively reconstruct the missing bio-chemical and bio-mechanical contexts of somitogenesis. Particularly, the µSDM focuses on modeling one important aspect of somitogenesis, somite boundary formation and associated morphogenetic cellular events, which eventually leads to delamination of new somites from the rostral end of the PSM. The µSDM effectively modularizes and thus is useful to decouple some critical molecular and cellular mediators of somitogenesis, such as external morphogen gradients and PSM tissue geometry. The modular bioengineering approaches utilized in the µSDM to decouple and independently control external biochemical gradients and tissue biomechanics are also useful for developing other controllable human embryo and organ models.

Other biomechanical aspects of somitogenesis, such as caudal elongation of the PSM, are not recapitulated in current µSDM. Even though HES7 oscillation dynamics along the R-C axis of the µSDM is shown, we are unable to explicitly correlate the segmentation clock with somite boundary formation dynamics in the temporal domain due to imaging limitations. It remains a future goal to apply the µSDM, in conjunction with high-resolution 4D imaging tools and signaling activity reporter lines, to study the interconnection between RA, FGF and WNT pathways and the segmentation clock and how such interactions regulate somite formation at both molecular and cellular scales.

Another important aspect about somitogenesis *in vivo* is the dynamic nature of morphogen gradients. As somites form and the PSM grows, FGF and WNT gradients shift caudally. Such dynamic signal gradients are not implemented in the current µSDM protocol but can be incorporated by adjusting morphogen dosages in medium reservoirs over time. As such, different rates and patterns of morphogen gradient shifting can be examined, which should help reveal the dynamic signaling interactions during somite development.

Despite its simplistic nature, the theoretical model we report here for somite size regulation based on a fracture mechanics-based framework can properly explain a primary correlation between somite and PSM length scales, thus supporting a pivotal role of mechanics in regulating somite boundary formation. Nevertheless, our current model is unable to recapitulate some occasional asynchronies between peak PSM length and peak somite size^27,28^, which suggests possible existence of secondary scaling. Mechanical gradients along the PSM and viscoelastic properties of mesodermal cells^35^ might need to be considered to fully rationalize both long-range interactions among somites, PSM and neighboring tissues, and local cellular activities in the forming somite and rostral PSM region.

In this study, we have developed a new hPSC-based, R-C patterned somite formation model. Compared with other existing, free floating culture-based, human somitogenesis models, exogenous morphogen gradients and spatial tissue confinements that somites and the PSM experience *in vivo* are explicitly introduced and integrated in the µSDM system. We further constructed a mechanical theory to explain the size dependency between somites and the PSM. By exploiting the compatibility of µSDM with live imaging and biomechanical and molecular characterizations and perturbations, we explored and validated the regulatory role of biomechanics in somite boundary formation dynamics. We envision that the µSDM will be useful for deconstructing the regulation and dysregulation of somitogenesis and ultimately promoting both fundamental knowledge and modeling of human skeletal and muscular deformities.

## AUTHOR CONTRIBUTIONS

Y.L. and J.F. conceived and initiated the project; Y.L. designed, performed, and quantified most experiments, including scRNA-seq data analysis and interpretation; Y.L. conducted theoretical modeling; Y.S.K. generated KO hPSC lines; X.X. developed MATLAB scripts for image processing and helped with scRNA-seq data analysis; N.K. and S.S. helped repeat experiments; O.P. provided HES7 reporter line; Q.Y. and O.P. helped with data interpretation and experimental designs; Y.L. and J.F. wrote manuscript. J.F. supervised the study. All authors edited and approved the manuscript.

## ACKNOWLEDGEMENTS

We thank Dr. Miki Ebisuya at TU Dresden for valuable discussions and Usha Kadiyala at Univ. of Michigan for assisting confocal imaging. This work is supported by Michigan-Cambridge Collaboration Initiative, University of Michigan Mcubed Fund, University of Michigan Mid-career Biosciences Faculty Achievement Recognition Award, National Science Foundation (PFI-TT 2213845, I-Corps 2112458 and CBET 1901718), and National Institutes of Health (R21 NS113518, R21 HD105126, R21 NS127983, R21 HD109635, R21 HD105192 and R01 NS129850). We also thank the Single Molecule Analysis in Real-Time (SMART) Center at Univ. of Michigan, seeded by NSF MRI-R2-ID award DBI-0959823 to Dr. Nils G. Walter, as well as Damon Hoff for training, technical advice and use of Atomic Force Microscopy. We acknowledge Michigan Medicine Microscopy Core for training and support in microscopy imaging, Michigan Advanced Genomics Core for scRNA-seq service, and Michigan Lurie Nanofabrication Facility for support in microfabrication.

## STAR METHODS

### RESOURCE AVAILABILITY

#### Lead contact

Further information and requests for resources and reagents should be directed to and will be fulfilled by the lead contact, Jianping Fu.

#### Materials availability

The cell lines generated in this study will be distributed upon request to other research investigators under a Material Transfer Agreement.

#### Data and code availability

R, Python, and MATLAB scripts used in this work are available from the lead contact upon reasonable request. Data supporting findings of this study are available within the article and its Supplementary Information files and from the corresponding authors upon request. ScRNA-seq data supporting this study’s results are deposited at Gene Expression Omnibus with accession number GSE236668.

### EXPERIMENTAL MODEL AND STUDY PARTICIPANT DETAILS

#### Cell lines

Human pluripotent stem cell (hPSC) lines used in this study include both human embryonic stem cells (H9, WA09, WiCell, NIH registration number: 0062; H1, WA01, WiCell, NIH registration number: 0043) and human induced pluripotent stem cells (NCRM1 hiPSC, a HES7-Achilles;pCAG-H2B-mCherry reporter^18^). H2B reporter and *TCF15*-knockout hPSC lines developed based on H9 hESCs are also used in this study. All protocols with hPSCs have been approved by the Human Pluripotent Stem Cell Research Oversight Committee at the University of Michigan, Ann Arbor. All hPSC lines have been authenticated by original sources as well as in-house by immunostaining for pluripotency markers and successful differentiation to the three germ layers. All hPSC lines are maintained in a feeder-free system for at least ten passages and authenticated as karyotypically normal. Karyotype analysis is performed by Cell Line Genetics. All hPSC lines are tested negative for mycoplasma contamination (LookOut Mycoplasma PCR Detection Kit, Sigma-Aldrich).

#### Cell culture

All hPSC lines are maintained in a standard feeder-free culture system using mTeSR medium (mTeSR; STEMCELL Technologies). H9 and H1 hESCs are cultured in tissue culture plates coated with lactate dehydrogenase-elevating virus (LDEV)-free, hESC-qualified reduced growth factor basement membrane matrix Geltrex (Thermo Fisher Scientific; derived from Engelbreth-Holm-Swarm tumors similarly to Matrigel). NCRM1 hiPSCs are cultured in tissue culture plates coated with hESC-qualified LDEV-free Matrigel (Thermo Fisher Scientific). Cell culture is visually examined during each passage to ensure absence of spontaneously differentiated, mesenchymal-like cells in culture. hPSCs between P50 and P70 are used for experiments.

#### Generation of H2B-eGFP hESCs

A CAG-H2B-eGFP H9 hESC line is generated as previously reported^5^. Specifically, H2B-eGFP (Addgene ID: 32610) is PCR amplified and cloned into an ePiggyBac vector with a constitutively active puromycin selection cassette^36^. The plasmid is co-transfected with pCAG-PBase (ePiggyBac transposase helper plasmid, provided by Dr. A.H. Brivanlou at Rockefeller Univ.) into H9 hESCs using Lipofectamine Stem (Thermo Fisher Scientific, STEM00003). Two days after transfection, CAG-H2B-eGFP H9 hESCs are selected with puromycin (1 µg mL^-1^; Thermo Fisher Scientific, A1113803) for 7 days.

#### Generation of *TCF15*-KO hESCs

*TCF15*-KO H9 hESCs are generated by targeting exon 1 of *TCF15* gene using CRISPR/Cas9-medited genome editing. Guide RNAs (gRNA) targeting the upstream and downstream introns spanning exon 1 of *TCF15* gene are designed using E-CRISP design tool (www.e-crisp.org/E-CRISP/designcrispr.html). The gRNAs are cloned into PX459-2A-Venus. The list of gRNAs is listed in **Key resources table**. The gRNAs are transfected into H9 hESCs and after 48 h, venus-positive cells are sorted by fluorescence-activated cell sorting (FACS). Sorted cells are plated on a tissue culture plate as single cells. After 7 days, individual hESC clones are isolated and are further genotyped for exon deletion by PCR using the primers listed in **Key resources table**.

### METHOD DETAILS

#### Microfluidic device fabrication

The microfluidic device for µSDM development consists of a polydimethylsiloxane (PDMS) structural layer attached to a PDMS micro-trench layer (**Supplementary Fig. 1a&b**). The PDMS structural layer is generated by mixing PDMS curing agent and base polymer (Sylgard 184; Dow Corning) at a ratio of 1:10 before casting PDMS prepolymer onto a microfabricated silicon mold and baking at 110°C for 1 h. Medium reservoirs (6 mm in diameter) and a loading port (1 mm in diameter) are then punched into the PDMS structural layer with Harris Uni-Core punch tools (Ted Pella).

To fabricate the micro-trench layer, PDMS molds are first generated by casting PDMS prepolymer with a 1:10 curing agent-to-base polymer ratio onto a microfabricated silicon mold and baking at 110°C for 1 h. Surfaces of PDMS molds are treated with air plasma for surface activation before silanization (Sigma-Aldrich, 448931-10G). The PDMS molds are then placed on PDMS prepolymer (1:20 curing agent-to-base polymer ratio) casted on a glass coverslip, before being baked at 110°C for 1 h. The PDMS micro-trench layers are then obtained after peeling off the molds.

Prior to experiments, micro-trenches are filled with 1% Geltrex (*v*/*v*) at 4°C overnight to coat their interior walls and bottoms. The PDMS micro-trench layers are then immersed in 1% bovine serum albumin (BSA; Thermo Fisher Scientific) solution at room temperature for 30 min. On Day 0, the PDMS structural layer is physically attached to the PDMS micro-trench layer, with micro-trenches visually aligned to be at the center of the microchannel in the PDMS structural layer.

The design of the PDMS structural layer includes three parallel channels that are partitioned by circular support posts (**Supplementary Fig. 1a&b**). The central channel in the PDMS structure layer is used for establishing chemical gradients through passive diffusion. Circular support posts separating the central channel from the other two channels are designed to constrain Geltrex solutions loaded into the central channel (see more information below) as well as to prevent air bubble trapping in the central channel during cell and Geltrex loading. The width and depth of micro-trenches are chosen to minimize air bubble trapping in the trenches during cell seeding, whereas the trench length is chosen to ensure sufficient difference in morphogen concentration along the trench length.

#### Development of µSDM

Between Day -2 and Day 0, colonies of hPSCs in tissue culture plates are treated with a basal medium supplemented with CHIR99021 (CHIR; 10 μM, STEMCELL Technologies) and LDN-193189 (LDN; 0.5 μM, STEMCELL Technologies), which is referred to as CL medium. The basal medium consists of Essential 6 (Gibco), GlutaMax (Gibco) and antibiotic/antimycotic (Gibco). On Day 0, cells in tissue culture plates are dissociated using Accutase (Sigma-Aldrich) at 37°C for 8 min before being suspended in DMEM/F12 (GIBCO) as single cells. Cells are then centrifuged and re-suspended in CL medium supplemented with Y27632 (10 μM, Tocris) at a density of 15 × 10^6^ cells mL^-1^. 10 μL cell suspension is then introduced into the central channel of the microfluidic device through its loading port on Day 0. Cells are allowed to settle into micro-trenches for 3 h, before the two medium reservoirs of the central channel are filled with CL medium supplemented with 10 μM Y27632. On Day 1, after aspirating culture medium from the central channel, 70% Geltrex (diluted in basal medium) is introduced into the central channel to establish a 3D culture environment. Starting from Day 1, the rostral reservoir connecting the central channel is filled with basal medium supplemented with retinoic acid (500 nM, STEMCELL Technologies) and PD173074 (400 nM, Tocris Bioscience), while the caudal reservoir is filled with CL medium supplemented with FGF8 (200 ng mL^-1^, PEPROTECH). Culture medium is then replenished daily.

#### Immunocytochemistry

To stain µSDM tissues, the PDMS structural layer is first removed from the microfluidic device prior to fixation of µSDM tissues. Cells and tissues are fixed in 4% paraformaldehyde (buffered in 1× PBS) for 12 h, and permeabilized in 0.1% SDS (sodium dodecyl sulfate, dissolved in PBS) solution at room temperature for 3 h. Samples are then blocked in 4% donkey serum (Sigma-Aldrich) at 4°C for 24 h, followed by incubation with primary antibody solutions at 4°C for 24 h. Samples are then labelled with donkey-raised secondary antibodies (1:400 dilution) at 4°C for 24 h. 4’,6-diamidino-2-phenylindole (DAPI; Thermo Fisher Scientific) is used for counterstaining cell nuclei. Both primary and secondary antibodies are prepared in 4% donkey serum supplemented with 0.1% NaN_3_. All primary antibodies used in this study are listed in **Key resources table**.

To clear µSDM tissues optically after immunofluorescence staining, µSDM tissues are incubated for 60 min in a refractive index (RI)-matching solution comprising 6.3 mL ddH_2_O (double distilled water), 9.2 mL OptiPrep Density Gradient Medium (MilliporeSigma), 4 g N-methyl-D-glucamine (MilliporeSigma), and 5 g diatrizoic acid (MilliporeSigma)^37^. For each µSDM sample, 50 μL of RI-matching solution is used.

#### Microscopy

Fluorescence imaging is conducted using an Olympus DSUIX81 spinning-disc confocal microscope. To image entire µSDM tissues, an array of partially overlapping images (50% overlap) are taken to cover entire µSDM tissues. Recorded images are stitched together using ImageJ plugin MIST. For *z*-stacking, images are acquired with a slice thickness of 0.5 μm. Low-magnification brightfield images are acquired using a Labomed TCM 400 inverted microscope equipped with a UCMOS eyepiece camera (Thermo Fisher Scientific). Brightfield live imaging is conducted using an inverted epifluorescence microscope (Zeiss Axio Observer Z1; Carl Zeiss MicroImaging) enclosed in an environmental incubator (XL S1 incubator, Carl Zeiss MicroImaging), maintaining cell culture at 37°C and 5% CO_2_. Fluorescence live imaging is conducted using an Olympus FV1200 confocal microscope equipped with a TOKAI HIT stage-top incubator to maintain cell culture at 37°C and 5% CO_2_. For z-stacking of fluorescence live imaging, images with a slice thickness of 5 μm are acquired.

#### Morphology quantifications

To quantify somite morphology, only rosette structures that show discernable outer and inner surfaces of the enveloping epithelium layer are selected. In ImageJ, an outline is manually drawn along the outer surface of rosettes, and morphological features such as area and circularity are automatically computed and extracted through the Measurement function of ImageJ.

#### RNA isolation and RT-qPCR analysis

On Day 5, the PDMS structural layer is first removed from the microfluidic device. µSDM tissues remaining on the PDMS micro-trench layer are cut into three even segments using a surgical scissor. RNA from each tissue segment is extracted using RNeasy mini kit (Qiagen) following the manufacturer’s instructions. A CFX Connect SYBR Green PCR Master Mix system (Bio-Rad) is used for RT-qPCR. An arbitrary Ct value of 40 is assigned to samples in which no expression is detected. Relative expression levels are determined as 2^−ΔΔCt^ with the corresponding s.e.m. Human GAPDH primer is used as endogenous control. All fold changes are defined relative to undifferentiated H9 hESCs. All analyses are performed with at least three biological replicates and two technical replicates. All primers are obtained from Ref^8,19,38^ and listed in **Key resources table**.

#### Single-cell dissociation and RNA-sequencing

To dissociate µSDM tissues into single cells, the PDMS structural layer is first removed from the microfluidic device, to expose and release µSDM tissues from micro-trenches. µSDM tissues are first cut into small pieces using a surgical knife and then incubated with Accutase for 2 - 3 h to obtain dissociated single cells. For scRNA-seq analysis of µSDM tissues at different time points, dissociated single cells from Day 2, 3, 4 and 6 µSDM tissues are harvested from 18 µSDM tissues. Dissociated single cells are collected into PBS containing 1% BSA before being centrifuged at 300 g for 5 min. Resultant cell pellets are re-suspended into single cells in PBS containing 1% BSA. Within 1 h after cell dissociation, cells are loaded into the 10× Genomics Chromium system (10× Genomics). 10× Genomics v.3 libraries are prepared according to the manufacturer’s instructions. Libraries are then sequenced using paired-end sequencing with a minimum coverage of 20,000 raw reads per cell using Illumina NovaSeq-6000. ScRNA-seq data are aligned and quantified using Cell Ranger Single-Cell Software Suite (v.3.1.0, 10× Genomics) against the Homo sapiens (human) genome assembly GRCh38.p13 from ENSEMBL.

#### Data integration, dimensionality reduction, and clustering

Analysis of scRNA-seq data and integration of scRNA-seq datasets are performed using R package Seurat (v.3.0.0.0, https://satijalab.org/seurat/)^39^. Default setups in Seurat are used unless noted otherwise. Briefly, each scRNA-seq dataset is filtered first based on the total number of genes detected and the percentage of total mitochondrial genes. Gene expression is then calculated by normalizing the raw count with the total count before multiplying by 10,000 and log-transformed. Top 2,000 highly variable genes are identified for each dataset using FindVariableFeatures. Datasets from different time points are then merged together. Cell cycle is regressed out based on cell cycle scores using CellCycleScoring during the data scaling process using SCTransform. PCA analysis (RunPCA) is then performed on filtered data followed by embedding into low dimensional space with Uniform Manifold Approximation and Projection (UMAP; RunUMAP) using dim 1:50, min.dist = 0.3, and n.neighbors = 5. Identification of cell clusters by a shared nearest neighbor (SNN) modularity optimization-based clustering algorithm is achieved using FindClusters with a resolution 0.2. To integrate multiple scRNA-seq datasets, count matrices of different datasets are first filtered and normalized separately before being integrated using IntegrateData. Integrated scRNA-seq dataset is analyzed following the standard Seurat pipeline. Annotation of cell clusters is based on expression of canonical lineage marker genes. The neural cluster identified is removed from further analysis. Differentially expressed genes (DEGs) are identified using FindAllMarkers, with min.pct = 0.1 and logfc.threshold = 0.25. Identified DEGs and their expression levels are summarized in **Supplementary Table 1**. Dot plots and feature plots are generated using DotPlot and FeaturePlot in Seurat, respectively. Heatmaps are plotted based on relative expression (*Z*-score) of top-20 gene signatures to distinguish each cell cluster under comparison.

#### RNA velocity analysis

FASTQ files generated by the Cell Ranger pipeline are used for RNA velocity analysis. Genome annotations GRCh38 are used for counting spliced and unspliced mRNA in each single cell. First, loompy fromfq is applied, with human genome assembly GRCh38 passed as an annotation, to generate the loom files containing both spliced and unspliced mRNA counts. Python package UniTVelo (v.0.2.2, https://unitvelo.readthedocs.io/en/latest/) is adopted to perform RNA velocity analysis^40^. Function ‘scv.pl.velocity_embedding_stream’ is used to project RNA velocities onto UMAP plots. All default parameters are used unless noted otherwise.

#### Trajectory inference and pseudotime analysis

R-package Slingshot is used for trajectory inference of the PSM-somitic cell lineage development^41^. Specifically, the merged dataset in Seurat is used as input to Slingshot. The rPSM cluster is assigned as the starting cell state. To visualize gene expression dynamics, expression levels of selected genes are first plotted along the pseudotime trajectory, and then fitted onto principal curves, which are further plotted as a function of pseudotime using plotSmoothers.

#### Comparison with monkey data

Three cell clusters (“PSM”, “Somitomere”, which is renamed in this study as “Nascent somite”, and “Early somite”) are chosen from the scRNA-seq dataset of a CS11 monkey embryo^16^. Monkey gene names are first projected to human ortholog gene names before the monkey dataset is integrated with the µSDM dataset using function IntegrateData with normalization.methd = “SCT”. UMAP embedding is then computed with first 30 principal components. Pearson’s correlations between cell clusters from the monkey CS11 dataset and the merged µSDM dataset are calculated using function cor.

#### Theoretical modeling of somite boundary formation

Given the physical similarity between somite segmentation and mechanical fracture, a fracture mechanics-based theory is developed to rationalize somite boundary formation process (**Fig. 4a**). In the model, the PSM is regarded as a homogeneous one-dimensional rod with length *L*, cross-sectional area *A* and *Young*’s modulus *E*. At the rostral end of the PSM tissue, a somite forming region with length *d* experiences an eigen-strain *ε*\* < 0, resulted from MET-induced cellular compaction. Specifically, a negative eigen-strain, describing an inelastic shrinking deformation, is resulted from tissue re-organization, which can persist under stress-free condition. Assuming that the PSM rod is fixed on both its ends, a simplification of the *in vivo* boundary condition resulted from tissues surrounding the PSM and somites, the strain energy in the entire PSM is calculated as *ψ^e^* (*d*) = *EA* (*ε*\*)^2^*d*^2^ / (2*L*). When a somite segmentation is initiated, a nascent somite delaminates from the rostral PSM region, releasing strain energy. However, this nascent somite formation leads to additional surface energy associated with newly generated somite and PSM interfaces as *ψ^s^* = 2*γA*, in which *γ* depicts surface energy density (surface tension). For somite boundary formation to initiate, the criterion of *ψ^e^* (*d*) ≥ *ψ^s^* needs to be satisfied, leading to a critical somite segment length *d* = [4*γL* / *E*(*ε*\*)^2^]^1/2^. Thus, our theoretical model predicts a scaling relation between somite length *d* and PSM tissue length *L* as *d* / *λ* ∼ (*L* / *λ*)^1/2^, where *λ* = 4*γ* / *E*(*ε*\*)^2^, and longer PSM tissues generate larger somites.

#### *In vivo* data extraction and model fitting

To validate the theoretical scaling law that correlates somite length scale with PSM length scale, data about somite and PSM sizes in zebrafish^28,30^, chicken^19,29^, and mouse^8,27^ embryo are extracted using software WebPlotDigitizer. For data in which PSM length and somite size are reported separately^27,30^, they are combined by correlating the associated developmental stages. Also, for the 1D-rod assumption in our theoretical model to hold for the PSM tissue, it’s necessary for the aspect ratio of the rod (or the PSM tissue) to be ≥ 5. Since the aspect ratio of somites *in vivo* is close to 1, in this work we only include data in which the PSM tissue length is at least five times of the nascent somite size.

#### *Young*’s modulus measurement by AFM

AFM force-distance (F-D) measurements are conducted using a TT-AFM (AFMWorkshop, South Carolina, USA) and AFM probes with a 2-µm diameter bead tip and manufacturer-calibrated spring constant of 0.064 N m^-1^ (NovaScan, Iowa, USA). Measurements are taken at 1 mm spacing along the length of µSDM tissues, with µSDM tissue samples moved between locations using a manual Vernier micrometer. Approximately 10 F-D curves are collected at each location along µSDM tissues, and those curves with effective loading are used for analysis. QPD sensitivity is determined in fluid by collecting F-D curves on a stiff PDMS surface. F-D curves are analyzed using AtomicJ^42^. Prior to the AFM measurements, the tissues are imaged with bright field microscope, and the regions without rosette formation is designated as the PSM regime.

#### Calculation and comparison of surface energy density

Model fitting using *in vivo* and *in vitro* data of somite and PSM sizes allows us to determine the value of *λ*. To examine the physiological relevance of *λ*, we can compare the surface energy value *γ* deduced from *λ* with the measured value from zebrafish mesoderm. By linearly interpolating AFM measurements along the tissue length, the average *Young*’s modulus *E* of the PSM tissue is about 0.74 kPa (**Fig. 4c**). The mean relative areal reduction of a forming somite recorded via live imaging is about 14.6% (**Fig. 4d**). Since we only consider tissue shrinkage of a forming somite along the R-C axis, the associated eigen-strain can be approximated as *ε*\* ≈ - 14.6% / 2 = -7.3%. Given *λ* = 4.41 µm from data fitting in **Fig. 4b**, we could determine the value of the surface energy density *γ* as 4.4 pN µm^-1^.

In a recent work by Maître *et al.*^32^, surface energy of zebrafish mesoderm is deconstructed into three parts, including cortical tension on cell-medium interface *γ*_cm_, cortical tension on cell-cell interface *γ*_cc_, and adhesion energy on cell-cell interface *ω*. The total energy associated with formation of somite boundary with area *A* can thus be written as *ψ^s^* = 2*γA* = (2*γ*_cm_ - 2*γ*_cc_ - *ω*)*A* = 2*γ*_cm_(1 - *γ*_cc_ / *γ*_cm_ - *ω* / 2*γ*_cm_)*A*. Based on Maître *et al.*^32^, for zebrafish mesoderm, *γ*_cm_ = 50 pN µm^-1^, *γ*_cc_ / *γ*_cm_ = 0.65, and *ω* / 2*γ*_cm_ = -0.06, which produces an effective *γ* = 20.5 pN µm^-1^.

#### Mechanical stretching of µSDM

A custom-developed cell stretching device (CSD)^43^ is employed to stretch µSDM tissues along their R-C axis direction (**Supplementary Fig. 4a**). Specifically, before the microfluidic device assembly, the PDMS micro-trench layer is attached to the CSD through plasma treatments on both the bottom surface of the PDMS micro-trench layer and the top surface of the CSD. The PDMS structural layer is then attached to the PDMS micro-trench layer, and µSDM tissue culture protocols proceed in the same way as previously described. On Day 5, the PDMS structural layer is removed from the device, and a mechanical loading with a 4-h period is applied to µSDM tissues inside micro-trenches for 24 h (**Supplementary Fig. 4c&d**). During this tissue stretching period, µSDM tissues are cultured in basal medium. To apply mechanical stretching of µSDM tissues, a trapezoidal voltage wave generated by a wave generator is converted to trapezoidal wave of vacuum pressure through a vacuum regulator (SMC Pneumatics, ITV0090). The trapezoidal wave of vacuum pressure is then loaded into the CSD to achieve uniaxial and periodical stretching of µSDM tissues inside micro-trenches (**Supplementary Fig. 4b**). On Day 6, µSDM tissues are fixed and processed for imaging. When fluorescence imaging is finished, the entire CSD device is placed under a brightfield microscope while the same vacuum pressure loading is applied. By measuring lengthening of micro-trenches, mechanical strain of µSDM tissues under different vacuum pressures are determined (**Supplementary Fig. 4d**).

#### Drug inhibition assays

For drug inhibition assays to block µSDM development, ADH-1 (0.2 mg mL^-1^, AdooQ Bioscience), cytochalasin D (10 µM, Tocris), and Y27632 (10 µM, Tocris) are supplemented to both rostral and caudal reservoirs of the microfluidic device between Day 3 and 5. In blebbistatin assays, blebbistatin (10 µM, Sigma Aldrich) is supplemented to both reservoirs between Day 1 and 5. All µSDM tissue samples are fixed on Day 5. All small molecules used in this study are listed in **Key resources table**.

### QUANTIFICATION AND STATISTICAL ANALYSIS

Statistical analyses are performed with OriginPro version 2023b. The statistical analysis method for each experiment is specified in the figure legend. For quantification, samples with air bubble trapped in microfluidic device during cell or gel loading are excluded. Samples with sub-optimal cell deposition on Day 1 are also excluded. No similar platform has been previously reported, thus the criteria were established specifically for this platform. Samples were randomly allocated to control and different experimental groups. However, no particular randomization method was used in this work.

### SUPPLEMENTARY FIGURE LEGENDS

**Supplementary Figure 1. Culture protocol and characterization of microfluidic somite development model (µSDM). a**, Schematics of µSDM device fabrication and its culture protocol. See **Methods** for µSDM design considerations. **b**, Image showing an assembled µSDM device. **c**, (left, top) Characterization of molecular diffusion in µSDM device by supplementing fluorescent dextran to the caudal reservoir. (left, bottom) Representative stitched micrographs showing diffusion of fluorescent dextran inside the microfluidic channel in the caudal (C)-to-rostral (R) direction over time. (right) Plot of fluorescence intensity along the microfluidic channel length over time as indicated. Fluorescence intensity is averaged across the microfluidic channel width. A stabilized fluorescent gradient pattern was established in the microfluidic channel within about 36 h. **d**, Derivation of presomitic mesoderm (PSM) cells from hPSCs. (left) Schematics of PSM differentiation protocol. (middle) Representative immunostaining images showing PSM cells stained positive for HOXC9 and TBX6, but negative for HOXC10, PAX3 or SOX2. (right) Bar plot showing percentage of TBX6^+^ PSM cells. Data are plotted as the mean ± s.d., with *n* = 3. **e**, Brightfield live imaging to examine cellular dynamics and spontaneous somite formation at the rostral end of a micro-trench over time. A forming boundary on Day 4 is marked by a white box. **f**, Representative stitched confocal image showing consistent rosette propagation from rostral ends of a micro-trench array. **g**, Representative stitched confocal micrographs showing µSDM on Day 4 and Day 6 stained for PAX3 and TBX6. Cell nuclei were counterstained with DAPI. Zoom-in views of three boxed regions are shown at the bottom. **h**, Representative stitched confocal micrographs showing µSDM on Day 5 stained for HOXC9, HOXC10 and TBX6. Cell nuclei were counterstained with DAPI. **i**, Schematic showing dissection of µSDM on Day 5 using a surgical scissor into rostral (R), middle (M), and caudal (C) tissue segments of equal lengths for downstream RT-qPCR analysis. **j**, Bar plots showing normalized expression of different somite and PSM markers as indicated, as a function of the three segments of Day 5 µSDM. *n* = 3, with *n*_µSDM_ = 6. *P* values calculated from two-sample *t*-tests are indicated. Scale bars: 1 mm (**c**); 100 µm (**d** & **e**); 200 µm (**f**); 200 µm for full-tissue images and 50 µm for zoom-in images (**g**); 200 µm (**h**).

**Supplementary Figure 2. Development of µSDM using different conditions and cell/tissue dynamics during somite formation revealed by live imaging. a,** Representative confocal micrographs showing rostral and caudal ends of µSDM derived from H1 hESC and hiPSC lines on Day 5 stained for PAX3 and TBX6 as indicated. Cell nuclei were counterstained with DAPI. **b**, Bar plot showing spatial regimes of somite formation in µSDM on Day 5 as a function of H9 and H1 hESC lines and a hiPSC line. For H9, *n*_µSDM_ = 4; for H1 and hiPSC, *n*_µSDM_ = 3. Data are plotted as the mean ± s.d. **c**, Areas of individual somites in µSDM on Day 5 as a function of H9 and H1 hESC lines and a hiPSC line. For H9, *n*_somite_ = 86; for H1, *n*_somite_ = 36; for hiPSC, *n*_somite_ = 41. Boxes and bars indicate interquartile ranges and median values, respectively, and squares and error bars indicate the mean ± s.d. *P* values calculated from two-sample *t*-tests are indicated. **d**, (left) Representative brightfield micrographs showing micro-trenches filled with hPSCs on Day 1 under different cell seeding density conditions as indicated. (right) Representative confocal micrographs showing individual somites in µSDM on Day 5 stained with DAPI under different cell seeding density conditions as indicated. **e**, Bar plot showing spatial regimes of somite formation in µSDM on Day 5 as a function of cell seeding density. For 8 × 10^6^ cells / mL, *n*_µSDM_ = 9; for 15 × 10^6^ cells / mL, *n*_µSDM_ = 8; For 25 × 10^6^ cells / mL, *n*_µSDM_ = 8. Data are plotted as the mean ± s.d. *P* values calculated from two-sample *t*-tests are indicated. **f**, Areas of individual somites in µSDM on Day 5 as a function of cell seeding density. For all conditions, *n*_µSDM_ = 4. For 8 × 10^6^ cells / mL, *n*_somite_ = 80; for 15 × 10^6^ cells / mL, *n*_somite_ = 115; For 25 × 10^6^ cells / mL, *n*_somite_ = 106. Boxes and bars indicate interquartile ranges and median values, respectively, and squares and error bars indicate the mean ± s.d. *P* values calculated from two-sample *t*-tests are indicated. **g**, Representative stitched confocal micrographs showing µSDM on Day 5 developed in micro-trenches with different widths stained with DAPI as indicated. Zoom-in views of boxed regions are shown on the right. **h**, Bar plot showing spatial patterns of somite formation in µSDM on Day 5 as a function of micro-trench width. For all conditions, *n*_µSDM_ = 3. Data are plotted as the mean ± s.d. *P* values calculated from two-sample *t*-tests are indicated. **i**, **j**, Areas (**i**) and lateral width (**j**) of individual somites in µSDM on Day 5 as a function of micro-trench width. For all conditions, *n*_µSDM_ = 3. For micro-trench width of 200 µm, *n*_somite_ = 39; for micro-trench width of 300 µm, *n*_somite_ = 60; for micro-trench width of 400 µm, *n*_somite_ = 74. Boxes and bars indicate interquartile ranges and median values, respectively, and squares and error bars indicate the mean ± s.d. *P* values calculated from two-sample *t*-tests are indicated. **k**, Representative stitched confocal micrographs showing µSDM on Day 5 stained with DAPI. Different FGF8 conditions were used for µSDM development as indicated. Somite formation regions are marked by while bars. Scale bar, 200 µm. **l**, Bar plot showing spatial regimes of somite formation in µSDM on Day 5 as a function of FGF8 concentration in the caudal reservoir. *n*_µSDM_ = 7 (50 ng/mL) and *n*_µSDM_ = 8 (200 ng/mL). Data are plotted as the mean ± s.d. *P* values calculated from two-sample *t*-tests are indicated. **m**, Areas of individual somites in µSDM on Day 5 as a function of FGF8 dose in the caudal reservoir. For both conditions, *n*_µSDM_ = 3. For FGF8 dose of 50 ng/mL, *n*_somite_ = 105; for FGF8 dose of 200 ng/mL, *n*_somite_ = 73. Boxes and bars indicate interquartile ranges and median values, respectively, and squares and error bars indicate the mean ± s.d. *P* values calculated from two-sample *t*-tests are indicated. **n**, Brightfield imaging of the rostral region of a single µSDM between Day 4 and 6, revealing spontaneous somite formation, beginning rostrally and extending caudally. **o**, Spatiotemporal distributions of the area (top) and circularity (bottom) of individual somites in the same single µSDM between Day 4 and 6 based on brightfield imaging. Each data point is from a single somite. **p**, Dynamic evolvements of area (top) and circularity (bottom) of individual somites between Day 4 and 6. Each light grey line represents an individual somite while black line shows the mean value. *n*_somite_ = 12. **q**, Confocal imaging of a H2B reporter line revealing sagittal view of cell dynamics in a forming somite close to the rosette front, which shows cellular compaction and reorganization, leading to the formation of a somite with an epithelial appearance and a closely packed circumferential ring of columnar-shaped cells, elongated in the radial direction. The rostral and caudal ends of the forming somite are marked by dashed lines. **r**, Confocal imaging of growth dynamics of a newly formed somite, showing dynamic cell movements and division within the somite epithelial ring. **s**, Confocal imaging of a formed somite showing centripetal movement of cells from the somite epithelium to the mesenchymal core of cells in the somitocoel. Red triangles mark movements of individual cells whereas blue triangles mark movements of cell clusters from the somite epithelium towards the somitocoel. Scale bars, 100 µm (**a**, **n**, **q** & **r**); 200 µm (brightfield images) and 100 µm (staining images) (**d**); 200 µm (full-tissue) and 100 µm (zoom-in) (**g**); 200 µm (**k**); and 10 µm (**s**).

**Supplementary Figure 3. scRNA-seq analysis of marker expression in µSDM. a**, (top) UMAP embedding of integrated single-cell transcriptome dataset of µSDM on Day 2, Day 3, Day 4 and Day 6, color-coded by cell identity annotation (top left) and pseudo-time (top right). (bottom) Feature plots showing expression patterns of key genes involved in the somitogenesis, including markers of the PSM, somite, sclerotome and dermomyotome as indicated. **b**, Heatmap of top-20 differentially expressed genes among all identified mesodermal lineages in µSDM. Color bar above the heat map indicates cell identity. **c & d**, Dot plots showing expression of *HOX*5-12 and RA and FGF signaling-related genes across different cell clusters in µSDM. Dot sizes and colors indicate proportions of cells expressing corresponding genes and their averaged scaled values of log-transformed expression, respectively. **e**, Expression levels of selected genes are fitted to principal curves to show general trends of their regulation. **f.** (top left) UMAP projection of integrated scRNA-seq dataset from µSDM and somitogenesis-related cells from a CS11 monkey embryo^16^. (top middle and right) UMAP projections of datasets from µSDM and the CS11 monkey embryo, separated from the integrated UMAP plot, with cell identity annotations indicated. (bottom) Feature plots comparing expression patterns of key PSM and somite markers in µSDM and the CS11 monkey embryo as indicated. cPSM, caudal PSM; rPSM, rostral PSM; N-SM, nascent somite; E-SM, early somite; SM, somite.

**Supplementary Figure 4. Mechanical stretching of µSDM. a**, (left) Experimental setup of a uniaxial cell stretching device (CSD) with a circular viewing aperture surrounded by a vacuum chamber. Two identical PDMS supports inserted symmetrically in the vacuum chamber divide the chamber into two identical vacuum compartments. (right) A bipolar suction generated by vacuum creates a uniaxial stretching field in the central region of a PDMS basal membrane, on which micro-trenches and µSDM are integrated for dynamic mechanical stretching. Please note that the length of µSDM (the R-C axis) is aligned to be parallel to the uniaxial stretching field. **b**, Brightfield images showing micro-trenches before and under mechanical stretching. Scale bar, 1 mm. **c**, Trapezoidal wave of vacuum pressure with a period of 4 h used for inducing µSDM stretching. **d**, Protocol for µSDM stretching experiments. µSDM tissues are maintained in basal medium and mechanically stretched between Day 5 and 6 for a period of 24 h, before being fixed and stained. After fluorescent imaging, µSDM tissues were stretched again under a brightfield microscope to measure applied strains. **e**, Schematic representation of generation of *TCF15*-KO hESCs using CRISPR/Cas9 genome editing. (left top) Targeted exon 1 (E1) of *TCF15* is shown in red and untargeted exon 2 (E2) is in black. gRNAs are designed to target the promoter region of *TCF15* and downstream intron of exon 1 as indicated. (left bottom) Sequencing results of control and *TCF15*-KO hESCs. (right) PCR validation of gene deletions.

### SUPPLEMENTARY VIDEOS

**Supplementary Video 1. Spontaneous somite formation in µSDM, beginning rostrally and extending caudally.** Time stamps indicate culture hours. Scale bar, 100 µm.

**Supplementary Video 2. Cell dynamics in a forming somite in µSDM.** Confocal imaging of a H2B reporter line shows cellular compaction and reorganization in a forming somite, leading to the formation of a somite with an epithelial appearance and a closely packed circumferential ring of columnar-shaped cells, elongated in the radial direction. Time stamps indicate culture hours. Scale bar, 100 µm.

**Supplementary Video 3. Growth dynamics of a newly formed somite in µSDM.** Confocal imaging of a newly formed somite shows a gradual increase of somite area, together with dynamic cell movements and division within the somite epithelial ring. Time stamps indicate culture hours. Scale bar, 100 µm.

**Supplementary Video 4. Dynamics of somitocoel cells in µSDM.** Confocal imaging of a formed somite showing centripetal movement of cells from the somite epithelium to the mesenchymal core of cells in the somitocoel. Time stamps indicate culture hours. Scale bar, 100 µm.

**Supplementary Video 5. Oscillation and traveling waves of *HES7* signals in µSDM.** Confocal imaging of a *HES7* reporter line shows oscillation and traveling waves of HES7 signals along the R-C axis of µSDM. Time stamps indicate culture hours. Scale bar, 200 µm.

### SUPPLEMENTARY TABLES

**Supplementary Table 1. List of differentially expressed genes identified from scRNA-seq data of µSDM. Expression levels of the genes are used to generate the heatmap in Supplementary Fig. 3b.**

Supplementary Table 2. List of genes used for calculating correlations between cell clusters identified from µSDM and *cy* monkey embryo.

