## Supplementary figures and images for "A human pluripotent stem cell-based somitogenesis model using microfluidics"

### Supplementary Figures

#### Supplementary Figure 1

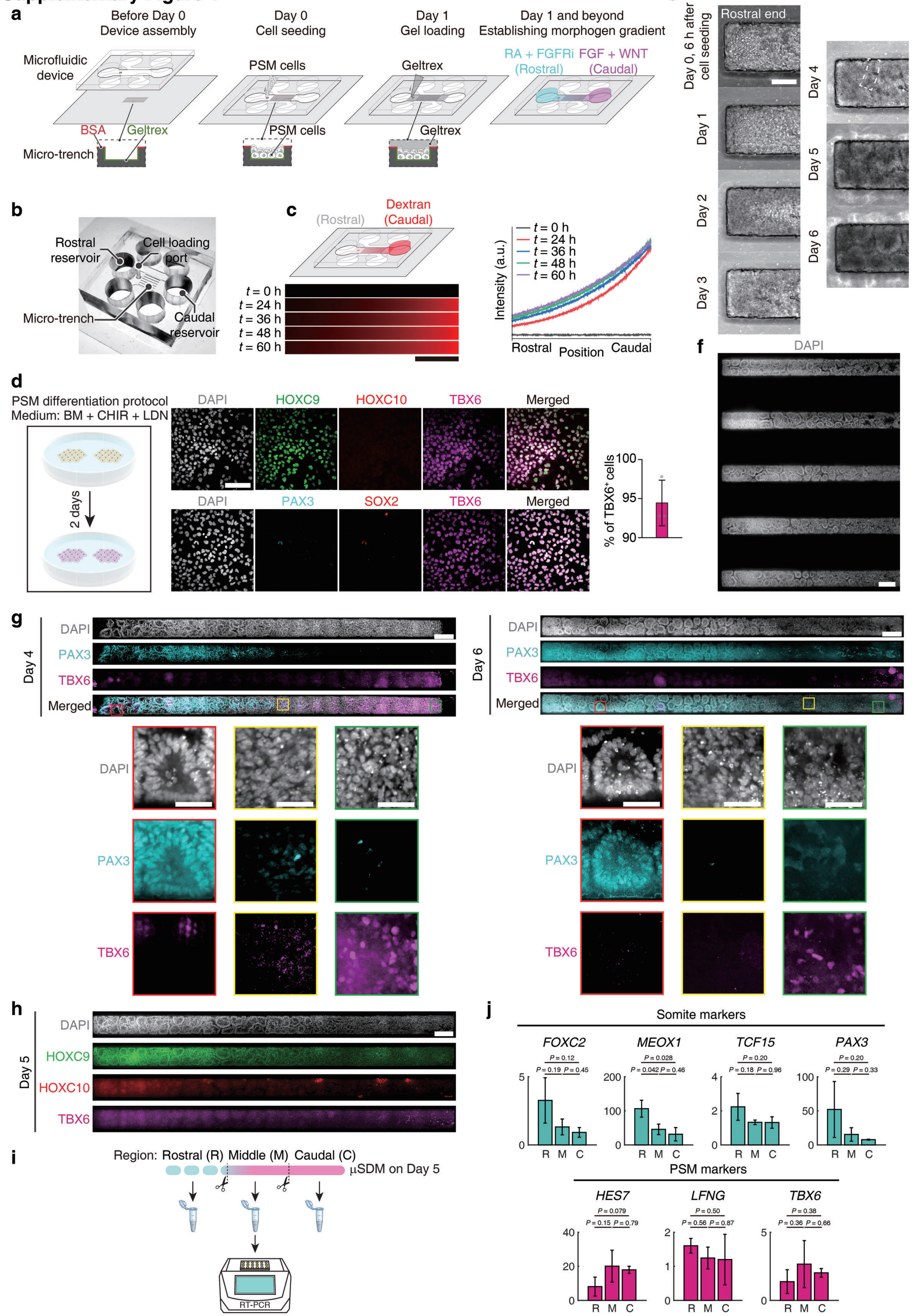

#### Supplementary Figure 2

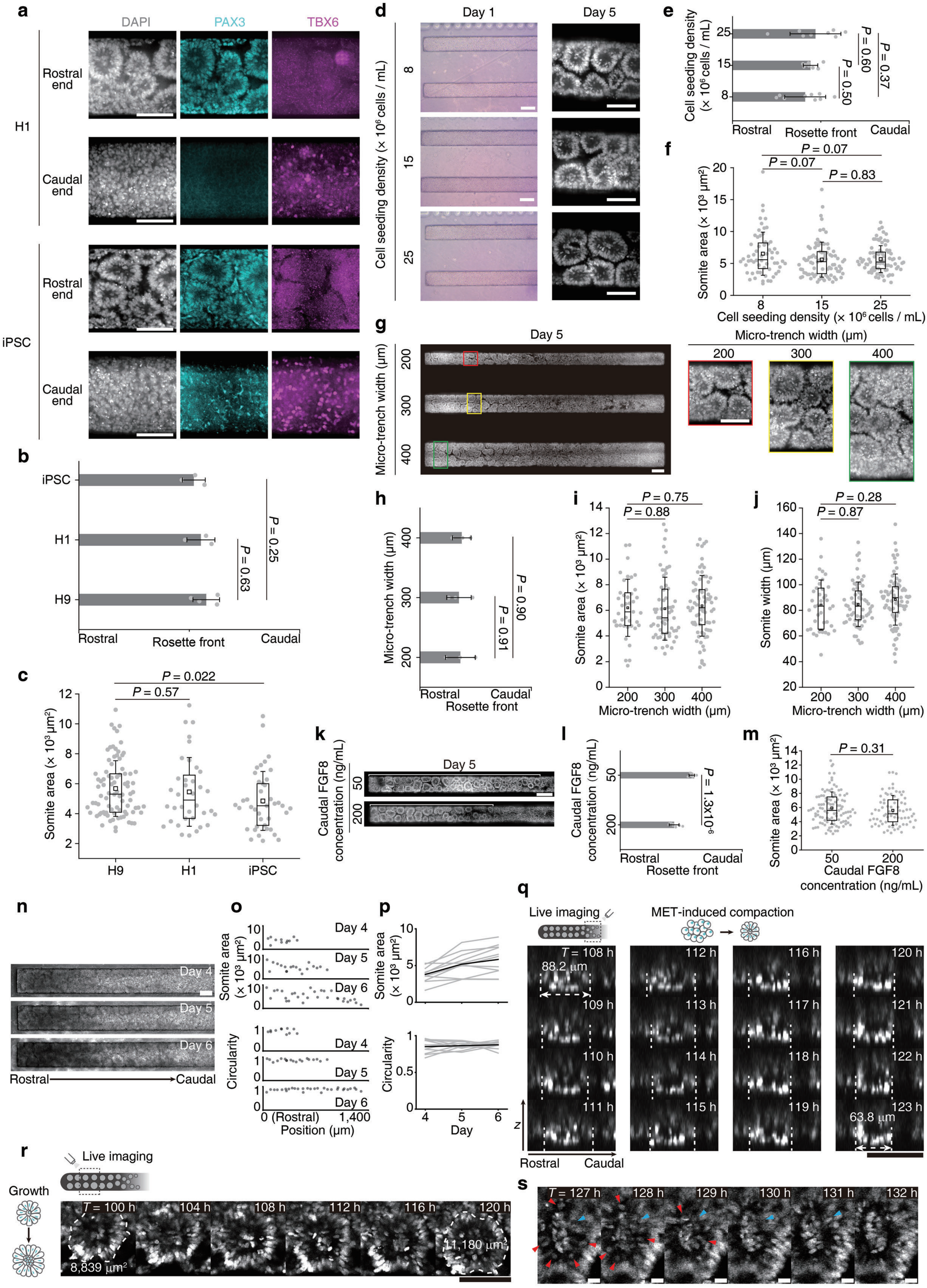

##### Supplementary Figure 3

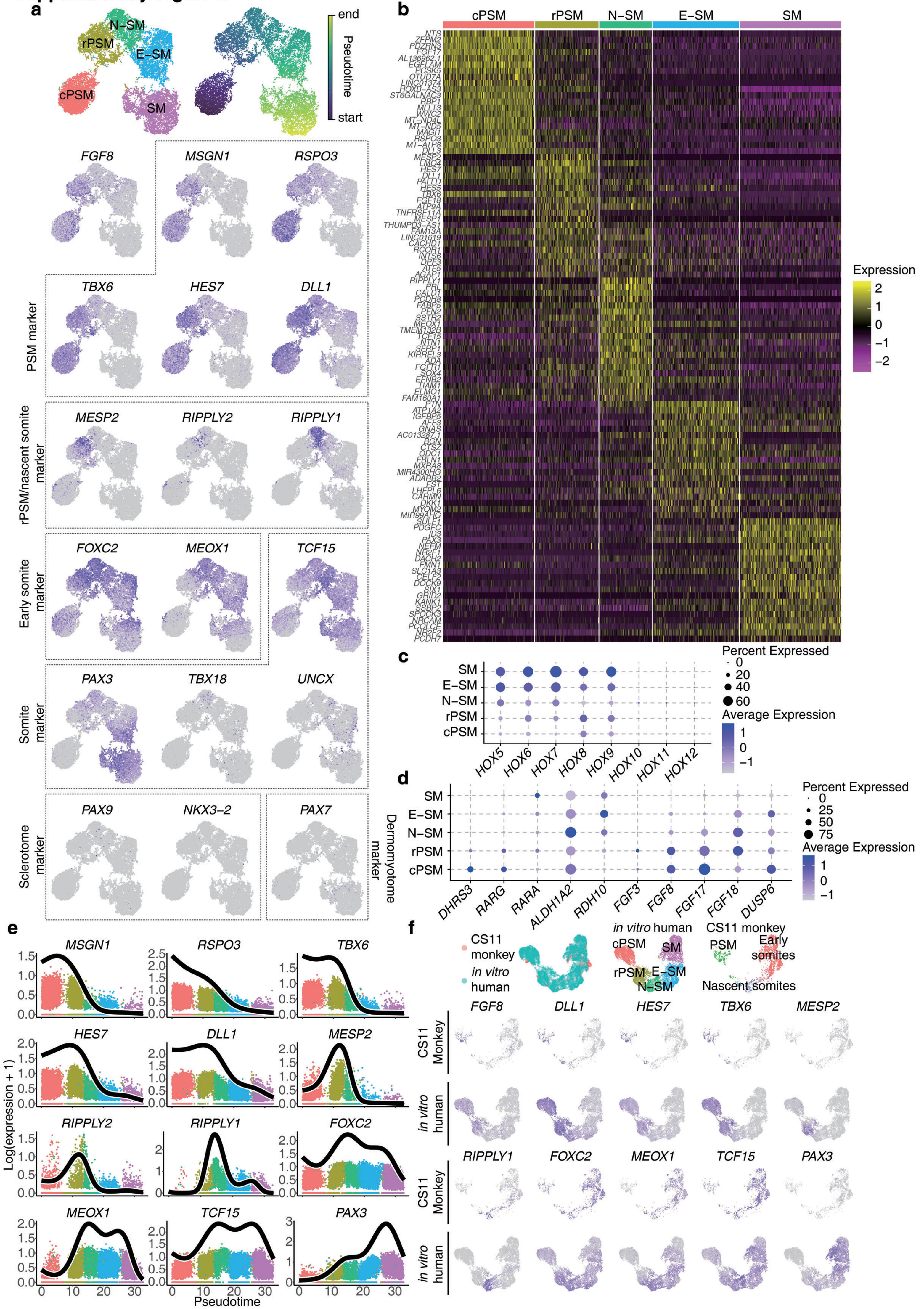

##### Supplementary Figure 4

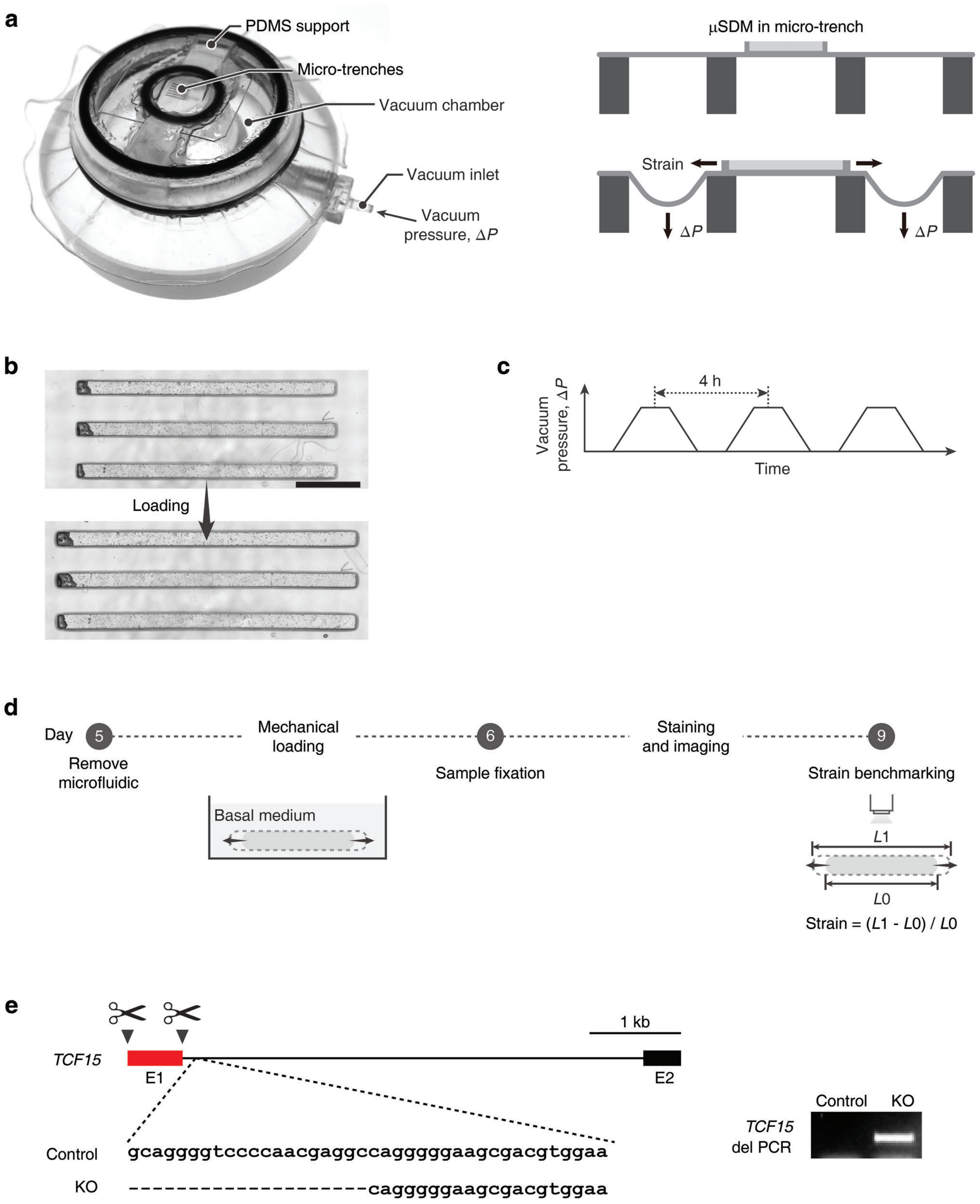
